# Isolated Nuclei Stiffen in Response to Low Intensity Vibration

**DOI:** 10.1101/2020.04.09.034546

**Authors:** J Newberg, J Schimpf, K Woods, S Loisate, P H Davis, G Uzer

**Affiliations:** Mechanical and Biomedical Engineering, Boise State University; Micron School of Material Science, Boise State University

**Author notes:** **Corresponding Author:** Gunes Uzer PhD, Boise State University, Department of Mechanical & Biomedical Engineering, 1910 University Drive, MS-2085, Boise, ID 83725-2085, **Email:**.

**Keywords:** Mesenchymal Stem Cells, Nucleus, LINC complex, Low Intensity Vibration

## Abstract

The nucleus, central to all cellular activity, relies on both direct mechanical input and its molecular transducers to sense and respond to external stimuli. While it has been shown that isolated nuclei can adapt to applied force *ex vivo*, the mechanisms governing nuclear mechanoadaptation in response to physiologic forces *in vivo* remain unclear. To investigate nuclear mechanoadaptation in cells, we developed an atomic force microscopy (AFM) based procedure to probe live nuclei isolated from mesenchymal stem cells (MSCs) following the application of low intensity vibration (LIV) to determine whether nuclear stiffness increases as a result of LIV. Results indicated that isolated nuclei were, on average, 30% softer than nuclei tested within intact MSCs prior to LIV. When the nucleus was isolated following LIV (0.7g, 90Hz, 20min) applied four times (4x) separated by 1h intervals, stiffness of isolated nuclei increased 75% compared to non-LIV controls. LIV-induced nuclear stiffening required functional Linker of Nucleoskeleton and Cytoskeleton (LINC) complex, but was not accompanied by increased levels of the nuclear envelope proteins LaminA/C or Sun-2. While depleting LaminA/C or Sun-1&2 resulted in either a 47% or 39% *increased* heterochromatin to nuclear area ratio in isolated nuclei, the heterochromatin to nuclear area ratio was *decreased* by 25% in LIV-treated nuclei compared to controls, indicating LIV-induced changes in the chromatin structure. Overall, our findings indicate that increased apparent cell stiffness in response to exogenous mechanical challenge of MSCs in the form of LIV is in part retained by increased nuclear stiffness and changes in chromatin structure.

## Introduction

Influence of the mechanical environment is perhaps most evident in resident stem cells which need to remodel their respective tissues in response to changing tissue mechanics (Rando and Ambrosio, 2018). For example, mesenchymal stem cells (MSCs) that replace and regenerate bone, when seeded into substrates with increasing stiffness tend to differentiate into bone lineage (Sun et al., 2018) through regulation of actin cytoskeleton dynamics and intra-nuclear Lamina (Buxboim et al., 2014; Swift et al., 2013). While cell structural changes and signaling events that take place within the focal adhesions and cytoskeletal compartments in response to environmental mechanical challenges are well studied (Lessey et al., 2012) the changes that occur inside the nucleus in response to physiological forces are less understood.

Dynamic forces applied to the exterior of the cell propagate through the cell via focal adhesions and cytoskeletal components, and reach the structural components at the outer and inner membranes of the nucleus, regulating nuclear structure (Martins et al., 2012). The nucleus is integrated within the cell structure through direct connections with cytoskeletal elements (Lombardi et al., 2011) through the LINC complexes (Linker of Nucleoskeleton and Cytoskeleton) composed of Nesprin and Sun subunits (Crisp et al., 2006). When forces are applied directly to the nucleus through LINC connections *ex vivo*, resulting in large strain deformations, nuclei stiffen via tyrosine phosphorylation of emerin (Guilluy et al., 2014) suggesting the nucleus is an active contributor to mechanotransduction. The nucleus also undergo structural re-organization in response to F-actin contractility by recruiting LINC complexes to apical stress fibers, leading to the accumulation of nuclear lamina element LaminA/C as well as changes in chromatin density under these stress fibers (Versaevel et al., 2014). At the level of DNA, forces applied at the cell membrane propagate through the cytoskeletal actin framework to cause deformations in chromatin (Tajik et al., 2016) and alter heterochromatin dynamics (Le et al., 2016). Nuclear stiffness is also in-part managed by the same toolset, Nesprins are anchored to the inner nuclear membrane through Sun-1&2 proteins that directly interact with structural elements such as nuclear pore complexes and LaminA/C (Hodzic et al., 2004; Padmakumar et al., 2005). Both Sun-1&2 and LaminA/C, play key roles in the structural integrity of the cell nucleus (Neelam et al., 2015). Inside the nucleus, both LaminA/C and chromatin have been shown to play independent roles in regulating nuclear mechanics (Stephens et al., 2017).

The elastic modulus of a cell nucleus can also be a marker of cell health. For example, it has been reported that the nuclei of Hepatitis C-infected cells are significantly softer than healthy controls, which was paralleled by downregulation of LaminA/C paired with upregulation of β- actin (Balakrishnan et al., 2019). Likewise, breast cancer cells exhibit large decreases in nuclear stiffness relative to healthy controls, showing downregulation of both LaminA/C and Sun-1&2 (Matsumoto et al., 2015). Furthermore, coinciding research showed chromatin decondensation as an additional component in highly metastatic cancer cells (Khan et al., 2018).

The nuclear envelope is subject to F-actin generated tension through LINC complex connections (Arsenovic et al., 2016). While cyto-mechanical forces can be generated in a multitude of ways, our group has focused on low intensity vibrations (LIV). LIV is a mechanical regime modeled after physiological, high frequency muscle contractions (Ozcivici et al., 2010). In MSCs, LIV promotes proliferation and osteogenic differentiation (Uzer et al., 2013) as well as increasing cell contractility by promoting GTP-bound RhoA (Ras homolog family member A)(Uzer et al., 2015). While we have further reported that daily LIV application up to 14 days increases stiffness of F-actin struts and results in increased mRNA expression of the LINC-related genes Nesprin-1&2, Sun-1&2, and LaminA/C in MSCs (Pongkitwitoon et al., 2016), the role of short term acute LIV application on the nuclear structural properties remains unknown.

Therefore, in this study we utilized AFM (atomic force microscopy) based nanoindentation measurements, immunostaining, and quantification of nuclear structural proteins to probe nuclear mechanical properties and morphology in response to acute bouts of LIV. We hypothesized that application of LIV to MSCs will increase nuclear stiffness.

## Results

### Cytoskeletal tension alters nuclear shape

We first investigated nuclear shape before and after our nuclear isolation protocol. **Fig. 1a** shows intact MSC (top) and isolated nuclei (bottom) after Hoechst 33342 staining to label DNA. As shown in **Fig. 1a**, the areas of isolated nuclei were visibly smaller than that of nuclei inside intact MSCs. As shown in Figures **1b** & **S1**, DNA structure and LaminA/AC organization remained intact within the isolated nuclei. We next evaluated the relative circularity of intact MSC nuclei versus isolated nuclei by measuring their XY (top view), XZ (side view), and YZ (side view) areas and combining these three-dimensional measurements to evaluate overall sphericity (**Fig. 1c**).

**Figure 1.**
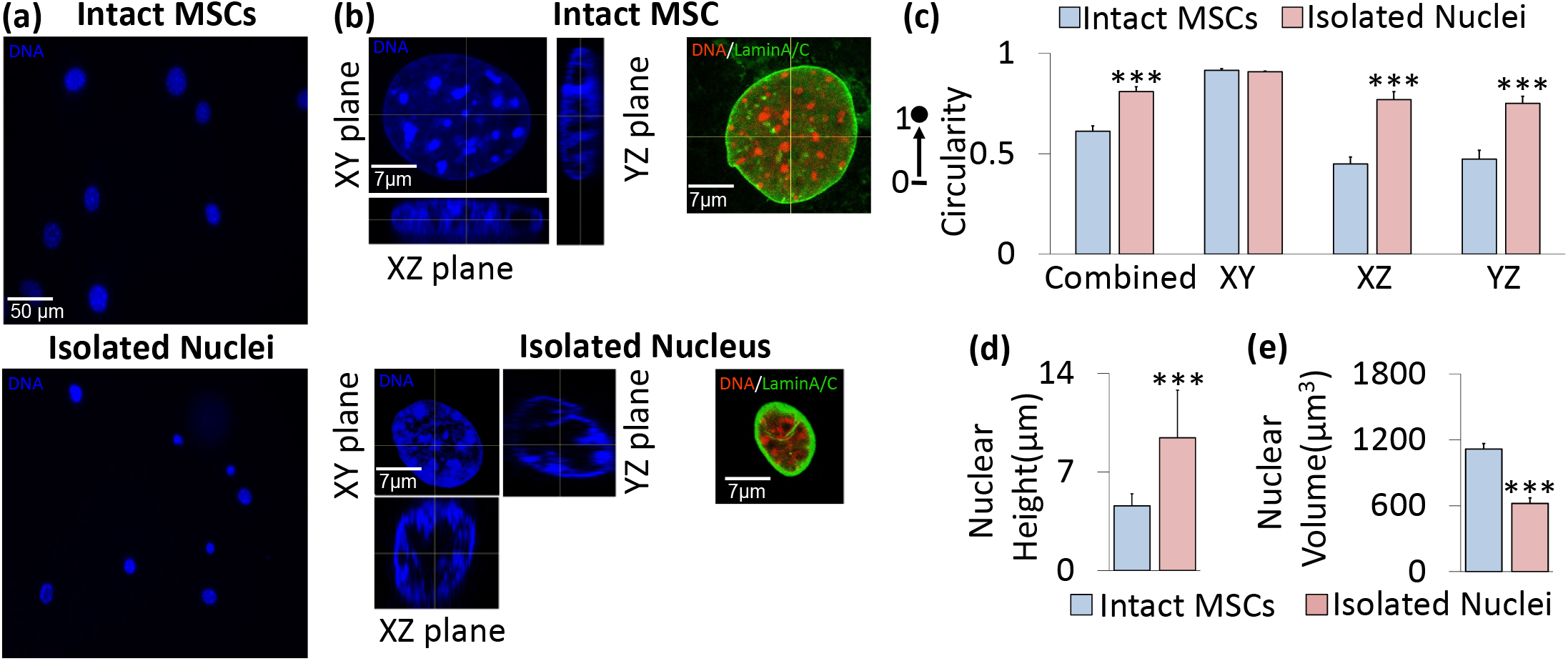
Nuclear geometry characteristics before and after nuclear isolation. **a)** Mesenchymal stem cells (top) and isolated nuclei (bottom) were stained with Hoechst 33342 and subsequently imaged for DNA using a fluorescence microscope. **b)** Confocal imaging of an intact nucleus (top) and isolated nucleus (bottom) with Hoechst 33342 fluorescence staining under 63x focus with 16 Z-stacks for each nucleus image. Representative intact MSC (top) and isolated nucleus (bottom) shown in XY, XZ, and YZ planes of focus. Staining against LaminA/C also indicated an intact nuclear lamina post-isolation. **c)** Shape profiles of the intact MSC nuclei versus isolated nuclei. The isolated and intact nuclei exhibited similar circularity in the XY plane (i.e., the horizontal plane of the cell culture dish or microscope slide), but showed significant differences in shape profiles in the vertical XZ and YZ planes (p<.001, N=10 isolated nuclei, N=10 intact nuclei), with isolated nuclei showing 52% and 45% higher circularity values in those planes, respectively. The combined data, which is an average of all three planes, shows a significant difference in sphericity (p<.001) between the isolated nuclei and intact nuclei, with the two types having sphericity values of 0.809 and 0.612, respectively. **d)** Average isolated nuclear height (9.43 μm) is approximately twice that of intact MSC nuclei (4.59 μm, N=10/group, p<.001). e) Volume decreases from 1,116 μm^3^ to 621 μm^3^ following isolation (N=10/group, p<.001. * p<.05, ** p<.01, *** p<.001, against control.

In the XY plane (i.e., the horizontal plane of the cell culture dish), intact and isolated nuclei showed no circularity differences. The isolated nuclei were 52% and 45% more circular than the nuclei of intact MSCs in the vertical XZ and YZ planes, respectively (p<.001). Combining the circularity values for each plane, the average circularity (i.e., sphericity) of the isolated nuclei was 0.809, which was significantly higher than that of the intact MSC nuclei (0.612, p<.001). Measures of nuclear height and volume (Figures **1d** & **1e**) showed that following nuclear isolation, height approximately doubled, increasing by 105% (p<.001), while overall volume decreased by 44% (p<.001).

### Nucleus significantly contributes to AFM-measured MSC stiffness

To investigate the mechanical properties of the nucleus, AFM-based nanoindentation was used to measure and compare elastic moduli values of intact MSC nuclei and isolated cell nuclei (**Fig.2a**). Prior to testing, tipless AFM MLCT-D probes were functionalized with 10 μm glass beads (Thermo Scientific 9010, Figures **2b** & **S2a**). Nuclei of intact MSCs and isolated nuclei were probed at the nuclear center visible in the XY plane (**Fig.2a**). To test whether fixation methods preserve nuclear stiffness, isolated nuclei were fixed in 2% paraformaldehyde (PFA) for 10 minutes and compared to untreated live controls. As shown in **Fig. 2c**, the results indicated a 4-fold increase in stiffness between PFA-treated and control nuclei (p<.01).

**Figure 2.**
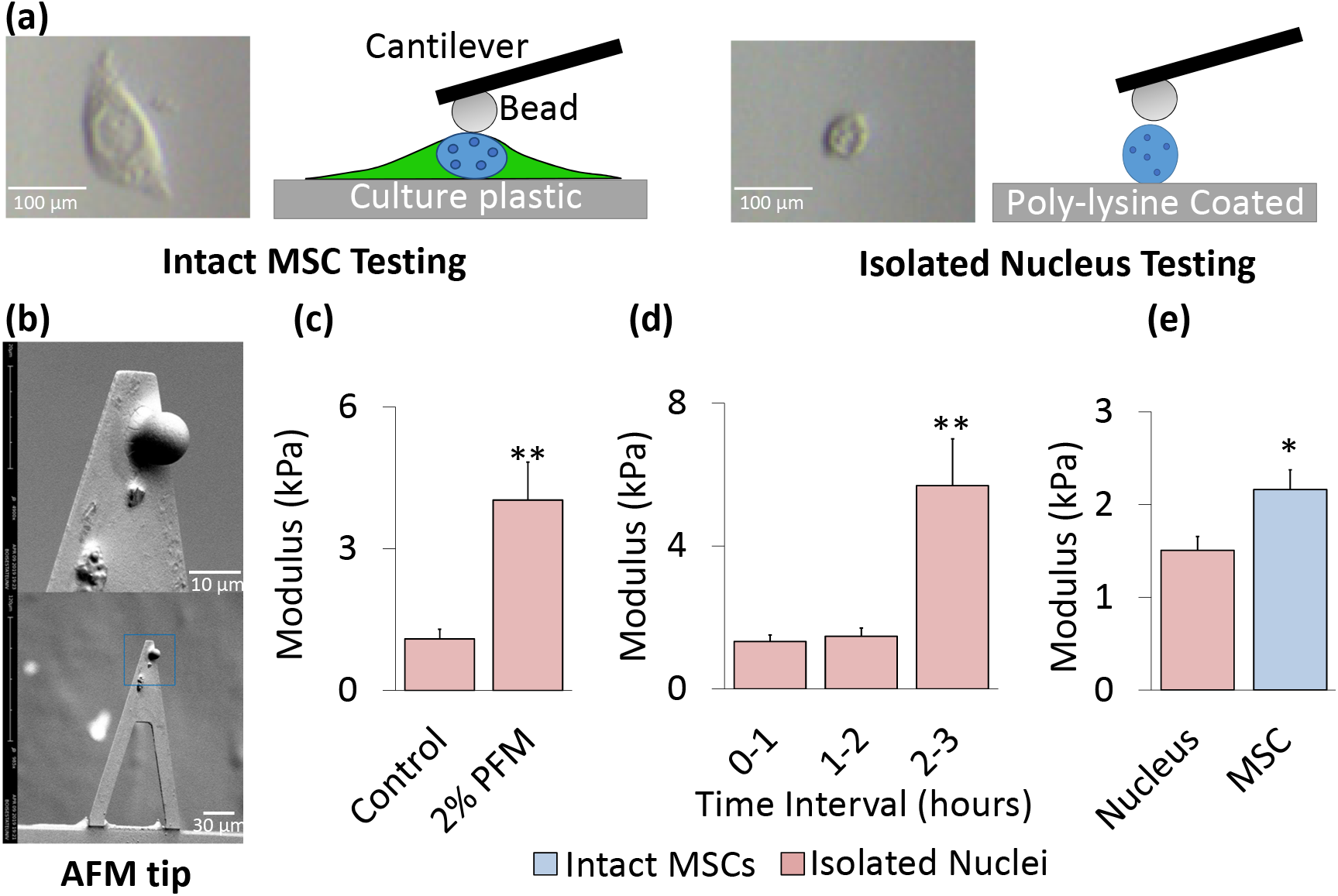
Nucleus significantly contributes to AFM-measured MSC stiffness. **a)** Nuclear isolation protocol involved plating of nuclei onto a 35 mm cell culture dish coated in poly-L-lysine. Isolated nuclei were incubated for 25 minutes at 37°C with 1 mL 1X PBS to ensure adhesion to the substrate immediately prior to AFM testing for up to 1 hour. Shown are both cartoons and actual optical images of intact MSCs and isolated nuclei as seen in the AFM camera. **b)** SEM image of a Bruker MLCT-D tipless AFM cantilever with a 10 μm diameter glass bead standard (Thermo Scientific 9010) attached. Scale bars were 30 μm (bottom image) and 10 μm (top image). **c)** Fixation of isolated nuclei in 2% paraformaldehyde (2% PFA) for 10 minutes showed an almost 4-fold increase in modulus compared to live controls (p<.01, N=10/grp). **d)** AFM measurements show no change in the elastic modulus of isolated nuclei over a 2-hour testing interval, after 2-hour window there was a 405% increase in the nuclear stiffness (p<.01). **e)** Isolated nuclei were identified via AFM and tested in the center of the nucleus to collect three individual measurements per sample. Intact MSCs and Isolated nuclei (both live) were plated on 35 mm dishes and tested for stiffness via AFM for one-hour intervals. The cell nucleus accounts for 69.7% of the total measured MSC stiffness with a modulus of 1.51 kPa (N=119). For comparison, the average stiffness of MSCs (passage 8-11) was 2.19 kPa (N=124). * p<.05, ** p<.01, *** p<.001, against control or against each other.

With fixation having been shown to significantly impact measured stiffness, we next determined the time window during which stiffness of live cell nuclei remains stable. Moduli of isolated live nuclei were measured across 3 one-hour time blocks, namely 0 to 1 hour, 1 to 2 hours, and 2 to 3 hours following isolation. **Fig. 2d** indicates that no change was observed in the mechanical properties of isolated nuclei up to two hours post-isolation, after which a large spike in stiffness occurred 2-3 hours following isolation (+406%, p<.01). Accordingly, a one hour post-isolation window for AFM-based nanoindentation testing was deemed safe and unlikely to skew the results. With this protocol established, we compared the stiffness of intact MSC nuclei (tested in the center of the nucleus) versus isolated nuclei within 1 hr of isolation. As shown in **Fig. 2e**, the average modulus of the isolated nuclei (1.5 kPa) was significantly softer than that obtained for the nuclei of intact MSCs (2.19 kPa, p<.05), indicating that the nucleus itself accounted for 69.7% of the overall MSC stiffness.

### Disruption of LaminA/C decreases nuclear stiffness and increases heterochromatin area in isolated nuclei

We next probed the effects of depleting known structural members in the nuclear membrane: Nuclear Lamina element, LaminA/C, and LINC complex elements Sun-1 and Sun-2 (**Fig. 3a**). Following siRNA depletion of LaminA/C (siLmn), co-depletion of Sun-1&2 (siSun), or control siRNA (siCntrl), groups were divided into either intact MSCs or underwent the nuclear isolation protocol. The siLmn group showed a 63.2% and 49.7% modulus decrease in intact MSCs (p<.05) and isolated nuclei (p<.01), respectively (**Fig. 3b**). Likewise, siSun treatment resulted in 50.7% and 27.7% decreased modulus for intact MSCs and isolated nuclei, respectively, but changes remained not significant compared to siCntrl (**Fig. 3b**).

**Figure 3.**
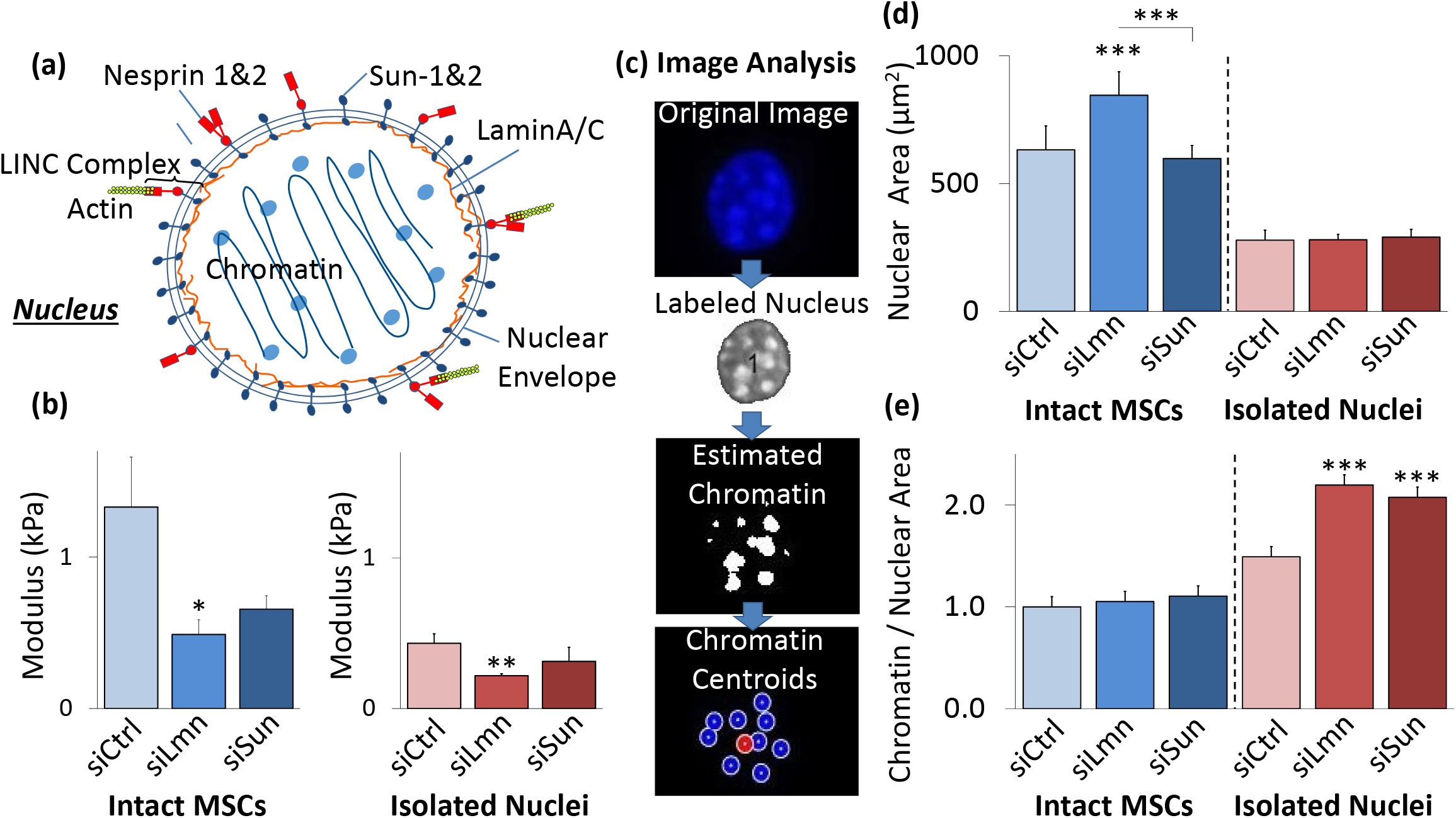
Disruption of LaminA/C decreases nuclear stiffness and changes structure. **a)** Schematic representation of the nucleus with nucleoskeletal and cytoskeletal connection components illustrated, including LaminA/C, Sun-1&2, Nesprin, and the KASH domain. **b)** SiRNA against nuclear LaminA/C (siLmn) and Sun-1&2 (siSun) decreased both isolated nuclear and intact MSC stiffness. Compared to siCtrl (N=21), siLmn and siSun treatments showed a significant decrease in the intact MSC modulus by 63.2% (p<.05, N=22) and 50.7% (p>.05, N=10), respectively. Likewise, when compared to siCtrl (N=21) the elastic modulus of isolated nuclei decreased by 49.7% (p<.01, N=21) and 27.7% (p>.05 N=10) in siLmn and siSun groups, respectively. **c)** A MATLAB code was constructed to evaluate differences in nuclear area and nucleoli size as determined by live Hoechst 33342 staining (see Supplementary Methods). **d)** Nuclear staining via Hoechst 33342 and epifluorescence imaging revealed that nuclear area increased by 33% within siLmn treated intact MSCs (p<.001, N=73), but not under Sun-1&2 depletion when compared to siCtrl (N=104). Nuclear area showed no significant changes within isolated nuclei control or experimental groups (N=245). **e)** Chromatin area to nuclear area ratios were calculated for all MSC and isolated nuclei for siCtrl, siLmn and siSun groups. Compared to controls (N=806), intact MSC did not show differences in chromatin to nuclear area ratio for LaminA/C (N=647) andSun-1&2 MSCs (N=489). In contrast, isolated nuclei showed significant increases in chromatin to nuclear area ratios for both LaminA/C (47.2%, p<.001, N=502) and Sun-1&2 (39.1%, p<.001, N=612) depleted nuclei compared to controls (N=416). * p<.05, ** p<.01, *** p<.001, against control or against each other.

Hoechst staining of DNA can be used to identify heterochromatin (Imai et al., 2017). We therefore quantified average heterochromatin size following LaminA/C or Sun-1&2 depletion using Hoechst 33342 and epifluorescence imaging (**Fig. 3c**). As shown in **Fig. 3d**, nuclear area increased 33% in siLmn MSCs (p<.001), but not in siSun MSCs. While overall nuclear area was smaller in isolated nuclei, no differences were observed in nuclear area between controls and isolated nuclei subjected to siRNA treatments (**Fig. 3d**). Combining the results presented in **Fig. 3c** and **3d**, the ratio of heterochromatin area to nuclear area was calculated to investigate possible changes in the heterochromatin condensation. As shown in **Fig. 3e**, measurement of the heterochromatin to nuclear area ratio in intact MSCs showed no changes between the control and siRNA-treated groups. In contrast, a comparison of the heterochromatin to nuclear area ratio of isolated nuclei showed significantly larger ratios compared to siCntrl for both the siLmn group (47.2% increase, p<.001) and the siSun group (39.1% increase p<.001).

### Low Intensity Vibration (LIV) stiffens MSCs and isolated nuclei

To evaluate nuclear mechanics following LIV, we subjected intact MSCs to either 2x or 4x low intensity vibration protocols. The LIV regimen is illustrated in **Fig. 4a**. As shown in **Figures 4b** and **4c**, following 2x LIV, intact MSCs showed a 71% increase in stiffness (p<.05), but there was no significant change in nuclear stiffness for isolated nuclei that underwent the same vibration protocol. In contrast, application of 4x LIV significantly increased the elastic modulus of both intact MSCs (419% increase, p<.001) and isolated nuclei (75%, p<.05).

**Figure 4.**
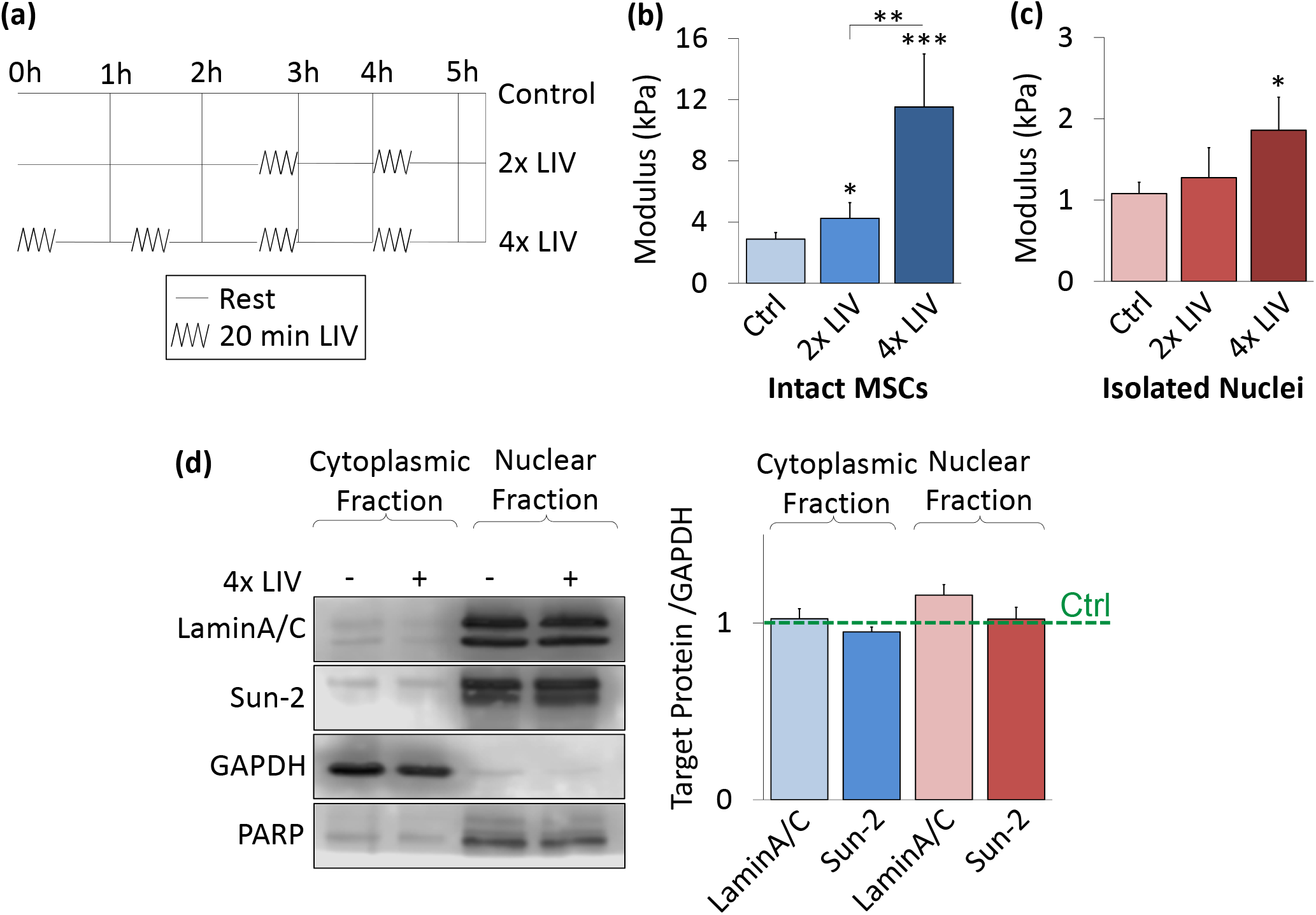
LIV stiffens intact MSCs and isolated nuclei. **a**) Low intensity vibration (0.7g, 90 Hz) was applied to MSCs at twenty-minute intervals with one-hour rest in between each vibration bout. 2x vibration included two, twenty-minute vibration periods, while 4x included four periods. **b)** In intact MSCs, 2x LIV increased AFM-measured elastic modulus on MSCs by 71% (N=31, p<.05). 4x LIV resulted in 419% increase in elastic modulus (N=33, p<.001) compared to control (N=32). **c)** Nuclear response to LIV was measured by applying the 2x LIV protocol to intact MSCs and then isolating nuclei to test stiffness. 2x LIV showed no significant increase in stiffness in comparison to control (N=20). Nuclei responded to 4x LIV by showing a 75% increase in modulus (N=24, p<0.05) when compared to control nuclei (N=27) post-LIV isolation. **d)** Western blotting for LaminA/C and Sun-2 show no changes in protein levels for 4x LIV groups compared to their respective controls (N=3/grp). * p<.05, ** p<.01, *** p<.001, against control or against each other.

To investigate the source of the observed changes in mechanical properties following LIV, the cytoplasmic and nuclear compartments were probed via western blotting. LaminA/C and Sun-2 protein levels following the 4x LIV protocol was quantified to test whether LIV results in upregulation of either protein. These protein levels were then compared to PARP and GAPDH as control markers. **Fig. 4d** shows no significant differences were observed in LaminA/C or Sun-2 protein levels, either in the cytoplasmic or nuclear fractions, following a 4x LIV protocol.

### Isolated nuclei maintain heterochromatin area after vibration

As LIV-induced stiffness increase was not accompanied by changes in LaminA/C or Sun-2 proteins, we probed possible changes in heterochromatin by comparing the heterochromatin to nuclear area ratio between 4x LIV and control samples for intact and isolated nuclei (**Fig 5a**). As shown in **Fig.5b**, intact MSCs subjected to 4x LIV showed no difference in nuclear area compared to controls. Likewise, there was no difference in nuclear area between isolated nuclei subjected to 4x LIV and the corresponding controls. As shown in **Fig.5c**, intact MSCs subject to 4xX LIV showed no difference in heterochromatin to nuclear area ratio compared to controls. In contrast to LaminA/C and Sun-1&2 depleted nuclei (**Fig.3e**) which showed higher chromatin to nuclear area ratio compared to control siRNA, isolated nuclei subjected to 4x LIV, showed a 25.4% decrease in chromatin to nuclear area ratio compared to non-LIV controls (p<.05).

**Figure 5.**
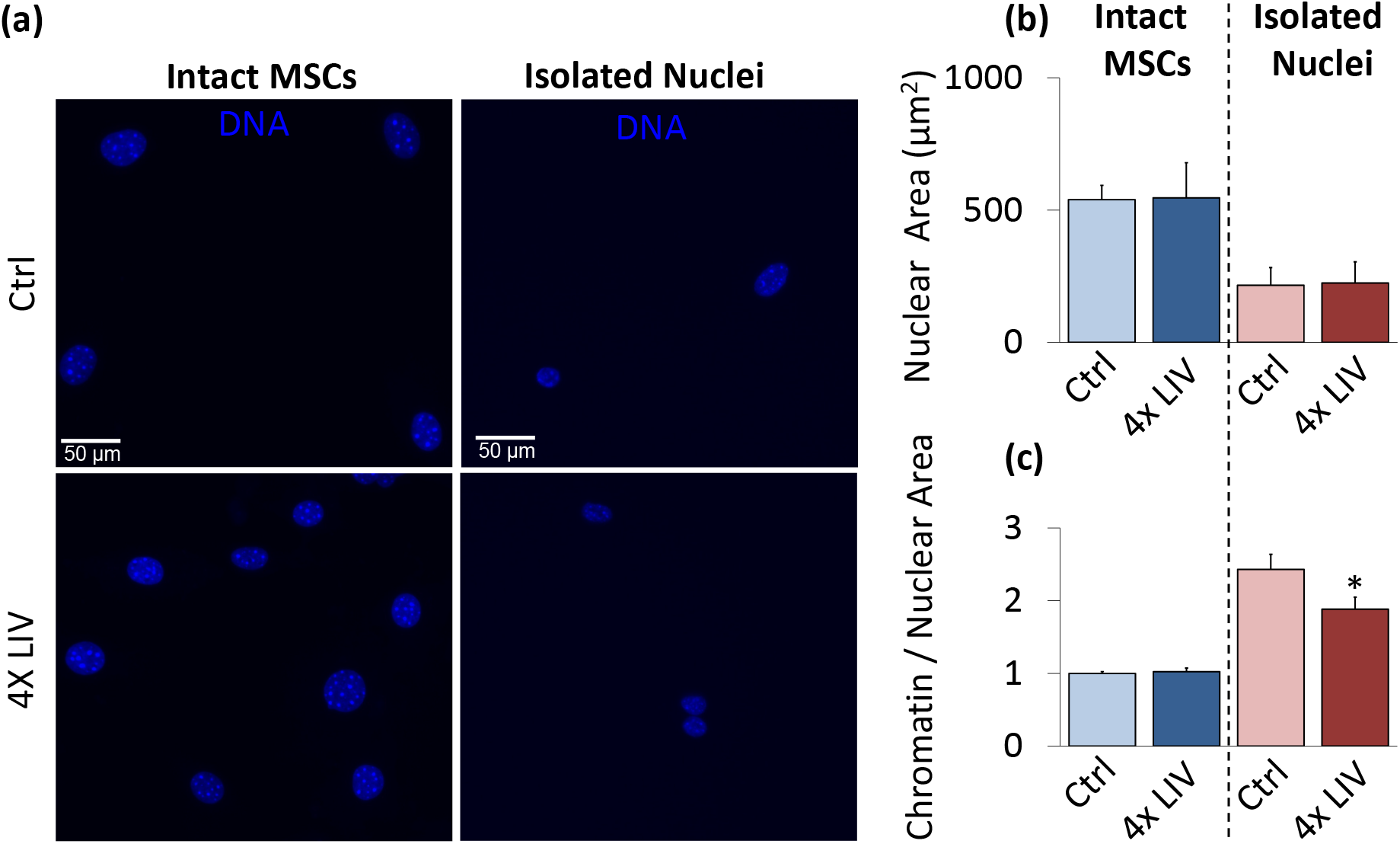
Isolated nuclei maintain heterochromatin area after vibration. **a)** Epi-fluorescence images of intact control MSC nuclei (upper left), intact 4x LIV MSC nuclei (lower left), isolated control nuclei (upper right), and isolated 4x LIV nuclei (lower right). Cells were treated with Hoechst 33342 to stain heterochromatin, fixed in 2% paraformaldehyde, and kept in PBS for imaging. **b)** Nuclear area was measured using MATLAB code to identify nuclear bounds under Hoechst 33342 staining. Intact MSCs subject to 4x LIV (N=216) showed no difference in nuclear area compared to controls (N=214). Likewise, there was no difference in nuclear area between control (N=90) and 4x LIV isolated nuclei (N=102). **c)** Chromatin measurements were analyzed via MATLAB analysis. Individual chromatin were averaged and compared to respective nuclear area. Intact MSCs subject to 4x LIV (N=2136) showed no difference in chromatin to nuclear area ratio compared to controls (N=2082). Isolated nuclei subject to 4x LIV showed a 25.4% lower chromatin to nuclear area ratio (N=480) than isolated controls (p<.05, N=360). * p<.05, ** p<.01, *** p<.001, against control.

### LINC function is required for LIV-induced nuclear stiffening

To test the requirement of cytoskeletal connections to the nucleus in the LIV response, we used LINC-compromised MSCs (referred to as +KASH). As depicted in **Fig.6a**, using previously established methods (Uzer et al., 2015), LINC complex function was disrupted by overexpressing a dominant negative form of the Nesprin KASH domain (pCDH-EF1-MSC1-puro-mCherry-Nesprin-1αKASH), which serves to disrupt the connection between Nesprin and Sun proteins via competitive binding to Sun proteins. Compared to empty control plasmid (pCDH-EF1-MSC1-puro-mCherry, referred to as mCherry), LINC-disrupted intact cells and isolated nuclei did not show any changes in stiffness (**Fig.S3**). As shown in Figures **6b** and **6c**, comparing intact MSC and isolated nuclear stiffness with or without 4x LIV showed no significant changes in elastic modulus between non-vibrated and vibrated MSCs with +KASH insertion.

**Figure 6.**
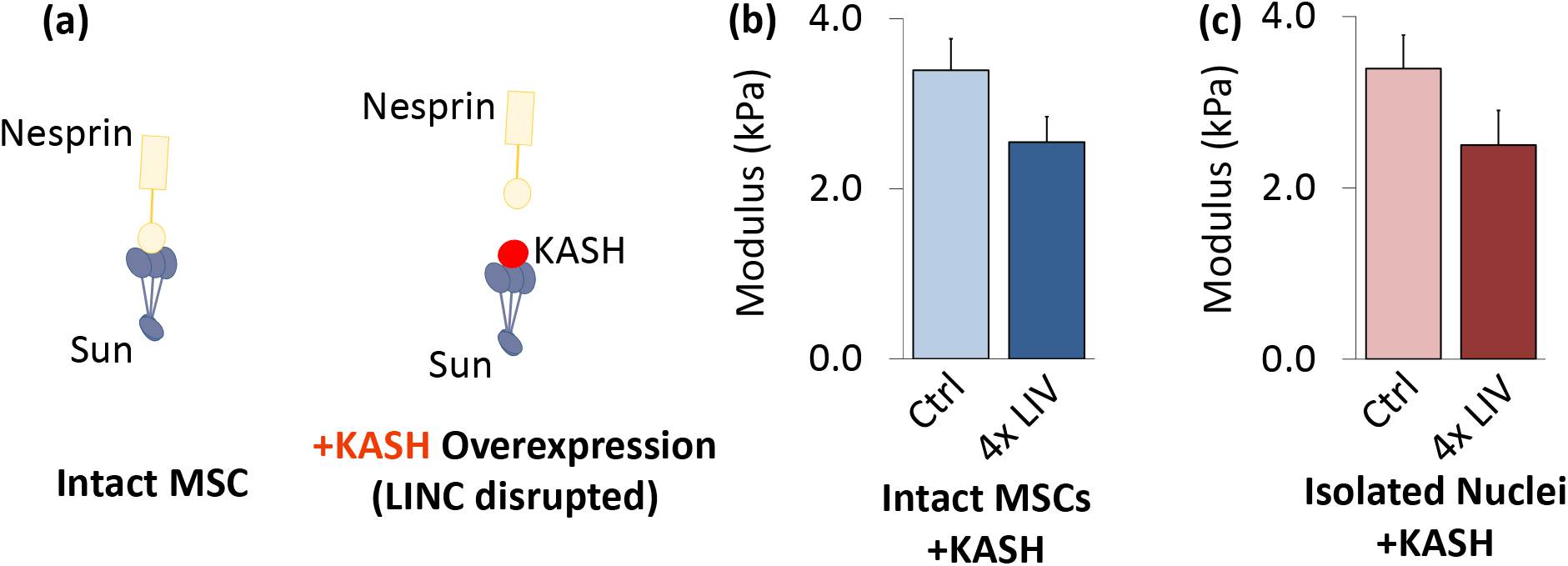
LINC function is required for LIV-induced nuclear stiffening. **a)** Overexpressing dominant negative form of Nesprin KASH domain was used to disrupt LINC function in MSCs. **b)** Intact MSCs and **c)** isolated nuclei did not show any response to 4x LIV (N=20/grp). * p<.05, ** p<.01, *** p<.001, against control.

## Discussion

Here we investigated the effects of a low magnitude, low intensity vibration (LIV) protocol on mesenchymal stem cell nuclei mechanical properties. Our results show that nucleus retains about 70% of its mechanical properties when isolated. Further the nucleus responds to low intensity vibration by increasing its apparent stiffness, not due to an increase in LaminA/C or Sun-2 protein amounts, but instead perhaps through changes in chromatin organization.

When cyto-mechanical forces that keep nucleus elongated was removed during nuclear isolation, isolated nuclei changed shape from 60% to 80% sphericity (**Fig. 1c**). We also observed a reduction in nuclear volume post isolation. While cytoskeletal force may, in part, account for the observed volume changes, nuclear volume depends on other factors including changes in pH, salt conditions, and temperature (Chan et al., 2017). Additionally, loss of protein structures during nuclear isolation has been reported (Paine et al., 1983). Although a potential limitation exist in the isolation technique, we confirmed changes in isolated nuclei stiffness via AFM-based measurements of elastic modulus (**Fig. 4c**).

When MSCs treated with siRNA protocols, both intact MSC and isolated nuclei modulus were consistently lower than non-siRNA counterparts (**Fig. 3b**), similar batch effects are also seen in KASH inserted MSCs **(Fig.6**). Increased stiffness of KASH inserted MSCs is probably due to extended culturing period during puromycin mediated selection process (see supplementary methods) as later passage MSCs were stiffer (**Fig. S4**) While this global shift in stiffness under siRNA treatment and plasmid overexpression remains as a limitation, in agreement with literature, depletion of LaminA/C showed significantly lower modulus compared to control siRNA (Lammerding et al., 2006). Softer nuclei, due to LaminA/C depletion also resulted in increased nuclear area (**Fig.3d**). This softening and concomitant increased nuclear area points to a weakening of nuclear structure. As expected, when Sun-1&2 were co-depleted, we did not observe any changes in the nuclear area of intact MSCs (**Fig.3d**), owing to an absence of cytoskeletal connections (Uzer et al., 2018). Following nuclear isolation (i.e. in the absence of cytoskeletal connections), both LaminA/C and Sun-1&2 depleted nuclei showed large increases in heterochromatin area. This observation suggests that in the absence of cytoskeletal constraints, nuclear Lamina and Sun proteins can have secondary effects on heterochromatin structure.

Acute application of LIV caused a significant increase in stiffness for both intact MSC nuclei and isolated nuclei (**Fig.4b**), suggesting that the changes in cellular stiffness and cytoskeletal tension (Pagnotti et al., 2019) in response to LIV were in-part retained by nucleus as increased stiffness even after nuclear isolation protocol. Though the stiffness was increased, there were no short-term changes in the levels LaminA/C and Sun-2 (**Fig.4c**). As the LIV-induced stiffening of isolated nuclei were lower than that of intact MSCs, our findings suggests other mechanisms responsible for MSC stiffening in response to LIV, such as the cytoskeletal connections to the nucleus. Additionally, disrupting the LINC complex connections and applying the same 4x LIV protocol showed no changes in elastic modulus. This further suggests that cytoskeletal components contribute to the overall stiffening response of both intact MSCs and isolated nuclei.

Image analysis indicated a retention of chromatin to nuclear area ratio in LIV treated MSCs post nuclear isolation (**Fig. 5c**). When cytoskeletal constraints were removed during nuclear isolation, nuclear to chromatin area ratio was increased in both control and LIV treated MSC nuclei, although nuclei subjected to 4x LIV showed significantly smaller, or more condensed heterochromatin compared to controls. These findings show that the effects of LIV were maintained by chromatin after the nucleus was mechanically separated from the cytoskeleton. While the mechanism by which this happens is again outside the scope of this current work, differences in DNA organization between isolated nuclei subject to LIV vs non-LIV counterparts may contribute to the increased stiffening of the nucleus (Stephens et al., 2018), suggesting that DNA organization plays a role in LIV-induced mechanoadaptation. Consequently LIV-induced decrease in the heterochromatin area post-isolation can potentially be a marker of altered gene access in response to environmental factors (Miroshnikova et al., 2017; Rubin et al., 2018).

Until now, it was previously unknown whether the nucleus responds to low intensity vibration in living cells. Our findings show that the nucleus is a mechanoresponsive element and responds to low intensity vibration by increasing its stiffness as confirmed by elastic modulus measurements. The mechanism(s) responsible for this stiffness regulation are not fully known, but our findings suggest that chromatin structure and cytoskeletal connections likely play a role in nuclear stiffening in response to LIV. Thus future studies aimed at understanding how LIV and other forms of mechanical signals regulate the mechanics of nucleoskeletal and chromatin structures may uncover new mechanisms by which forces regulate gene expression and cellular decisions.

## Materials and Methods

### Cell Culture

Primary mouse mesenchymal stem cells (MSCs) were extracted as previously described (Case et al., 2010; Peister et al., 2004). MSCs between passages seven (P8) and eleven (P11) were used during experiments for consistency, as higher passages showed significantly higher moduli, **Fig. S4**. For sub-culturing, cells were re-plated at a density of 1,800/cm^2^ and maintained in IMDM (12440053, GIBGO) supplemented with 10% FCS (S11950H, Atlanta Biologicals) and 1% Pen/Strep. For whole cell experiments, MSCs were plated into 35 mm diameter dishes prior to application of LIV. For nuclear extraction experiments, cells were maintained in 55 cm^2^ culture dishes until 80% confluency (approximately 1.5 – 2 million cells) prior to application of LIV. Transfections and siRNA were applied 72 hr prior to isolation protocols (see Supplementary Methods).

### Nuclear Isolation

MSCs were gently removed from plates by scraping in 9 mL of 1x PBS and centrifuged at 1100 hypotonic buffer (0.33 M sucrose, 10 mM HEPES, pH 7.4, 1 mM MgCl_2_, 0.5% w/v Saponin) and Centrifuge). For western blots, the cytoplasmic fraction (supernatant) and nuclei (pellet) were saved separately. For AFM experiments, the cytoplasmic supernatant was aspirated and nuclei were resuspended in 100 μL of hypotonic buffer (Buffer A). To gently separate cytoplasmic debris from nuclei, the resuspended pellet was added to 400 μL of Percoll (Sigma Aldrich) + were then plated in a 0.01% poly-L-lysine coated 35 mm cell culture dish and incubated for 25 minutes for adherence.

### Low Intensity Vibration (LIV) Protocol

Vibrations were applied to MSCs at a peak magnitudes of 0.7 g at 90 Hz for 20 minutes at room temperature(Sen et al., 2011). Controls were handled the same way, but the LIV device was not turned on. LIV was applied as either, two (2x) or four (4x), twenty-minute vibration periods with one-hour rest in-between.

### Atomic Force Microscopy (AFM)

Force-displacement curves for both intact MSC nuclei and isolated nuclei acquired using a Bruker Dimension FastScan AFM. Tipless MLCT-D probes (0.03 N/m spring constant) were functionalized with 10 μm diameter borosilicate glass beads (Thermo Scientific 9010, 10.0 ± 1.0 μm NIST-traceable 9000 Series Glass Particle Standards) prior to AFM experiments using UV-curable Norland Optical Adhesive 61 (**Fig. S2a**). To ensure accurate force measurements, each probe must be individually calibrated. Accordingly, a thermal tune was conducted on each probe immediately prior to use to determine its spring constant and deflection sensitivity (**Fig. S2b**). MSCs and nuclei were located using the AFM’s optical microscope and engaged on using a minimal force setpoint (2-3 nN) to minimize damage prior to testing. Ramps were performed over the approximate center of each nucleus for all samples. Force curve ramping was performed at a rate of 2 μm/sec over 2 μm total travel (1 μm approach, 1 μm retract; **Fig. S2c**). Three replicate force-displacement curves were acquired and saved for each nucleus tested, with at least 3 seconds of rest between conducting each test. Measurements that showed minimal contact with the nucleus were discarded and performed again to ensure an adequate depth of probing.

Measured force-displacement curves were analyzed assuming Hertzian (spherical contact) mechanics(Guo et al., 2012), employing Bruker’s Nanoscope Analysis software package to obtain the elastic modulus of the samples. Accordingly, MSC and nuclear stiffness were quantified in Nanoscope Analysis using a best-fit curve to a Hertzian model. The point of initial contact was visually selected, and the curve was analyzed until the *R*^2^ value was greater than 0.95 (p<0.05). The modulus of each individual sample (i.e., intact MSC or isolated nucleus) was reported as the average of three consecutive measurements. Group averages were then obtained from the average values for the individual samples.

### Statistical Analysis

Unless indicated otherwise in figure legends, results are expressed as mean ± standard error of the mean throughout. All experiments were replicated at least three times to ensure reproducibility. Densitometry and other analyses were performed on at least three independent experiments. As we previously reported (Uzer et. al 2018, Touchstone et al. 2019), for western blot data, differences between treatments within each biological replicate were assumed to follow a normal distribution due to large mean sample; thus for these comparisons, we have used two-tailed un-paired T-tests (Figure 4d). For other comparisons, we used a non-parametric two-tailed Mann-Whitney U-test (Figures 1, 2c, 2e, 5, 6, S3 and S4) or Kruskal-Wallis test followed by Tukey multiple comparison (Figures 2d, 3 and 4b). P-values of less than 0.05 were considered significant.

## APPENDIX: Supplementary Figure Legends & Tables and Methods

### Supplementary Figure Legends

**Figure S1.**
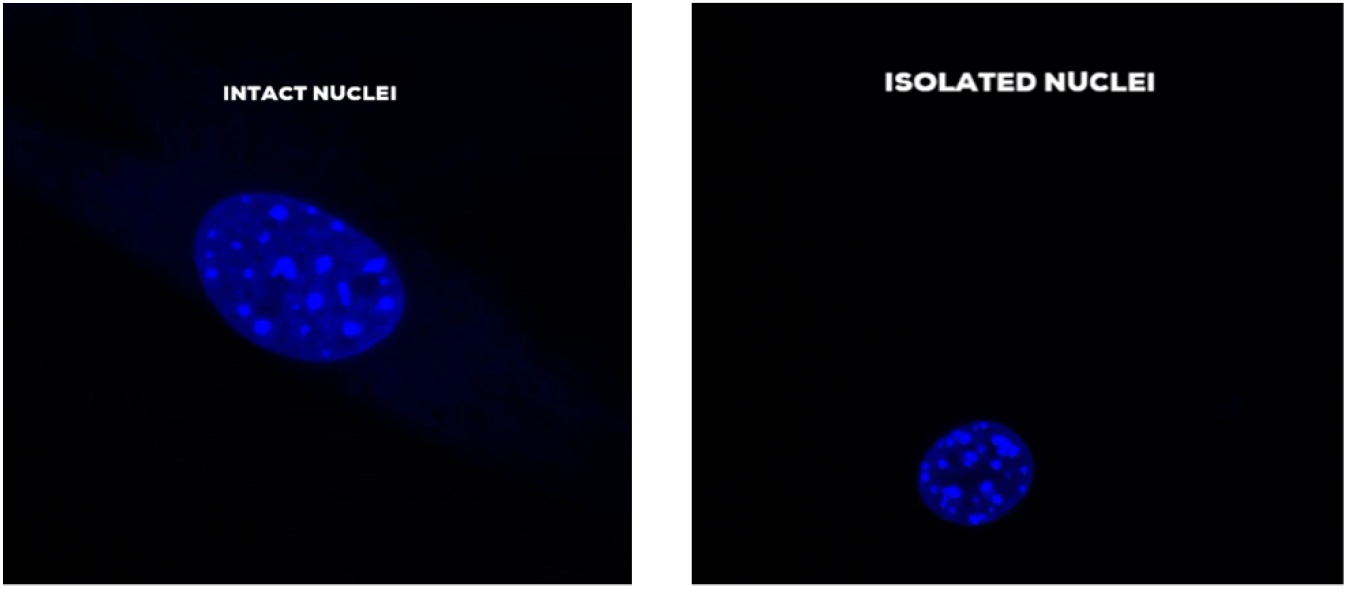
Nuclei were either left intact inside MSCs or isolated (isolation protocol, methods), stained with Hoechst 33342, and fixed in 2% paraformaldehyde. As shown in these videos, a Leica 6500 confocal microscope was then used to acquire vertical stacked images of both nuclei inside intact MSC samples (left) and isolated nuclei (right).

**Fig. S2.**
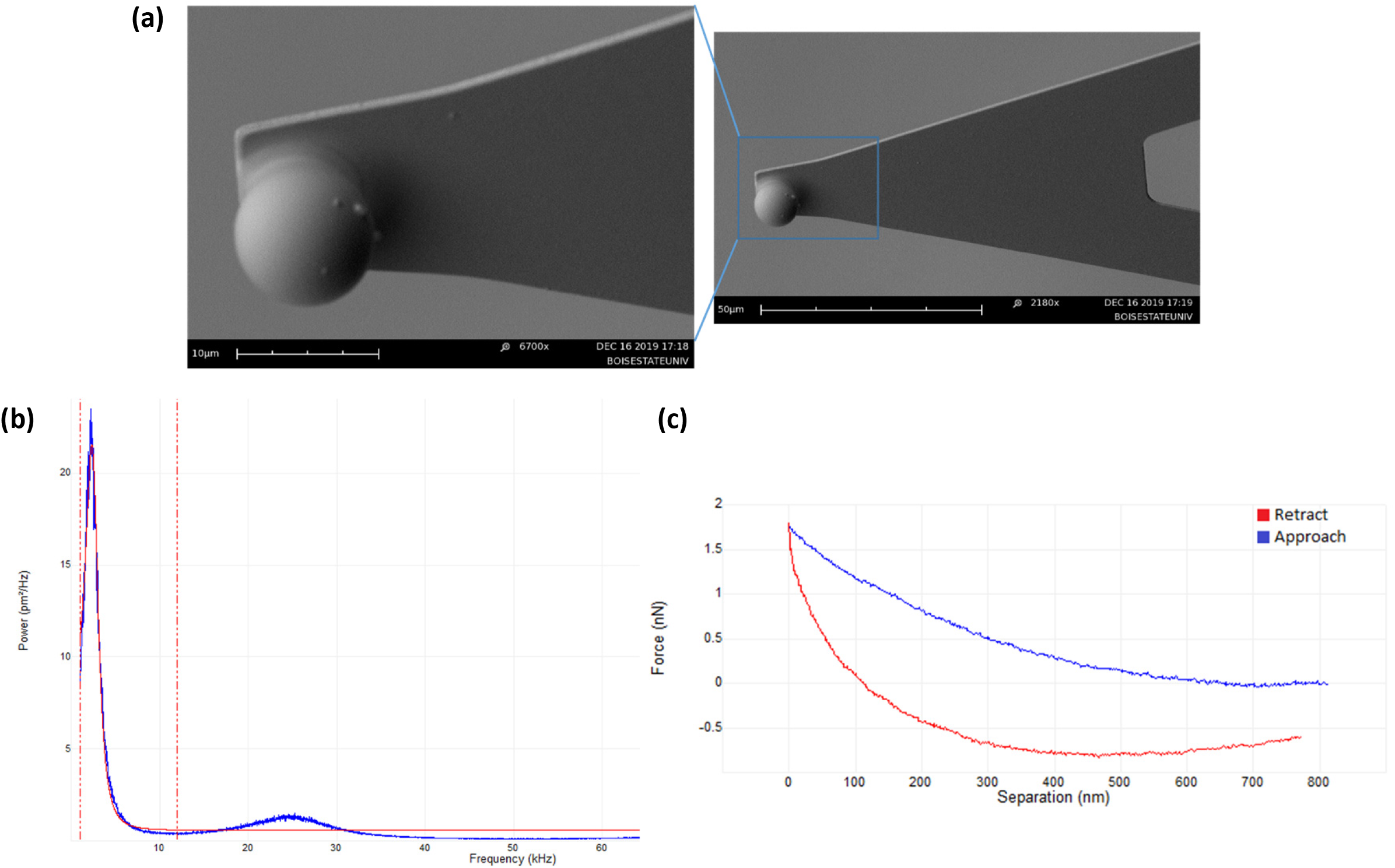
**a)** SEM image of a Bruker MLCT-D AFM probe functionalized with a 10 μm diameter glass bead (Thermo Scientific 9010 NIST-traceable 9000 Series Glass Particle Standard, 10.0 ± 1.0 μm). **b)** Representative thermal tune of an MLCT-D probe functionalized with such a 10 μm glass bead. Nominal resonance frequency of the MLCT-D cantilever prior to bead functionalization is 10-20 kHz, with the added weight of the bead causing the resonance to shift to lower frequency. A simple harmonic oscillator fit was used (red) to model the resonant response of the cantilever in a fluid environment. **c)** Representative ramp data. Ramping to a maximum excursion/extension of 1 μm was performed at a rate of 1 Hz (i.e., 2 μm/sec load/unload velocity).

**Fig S3.**
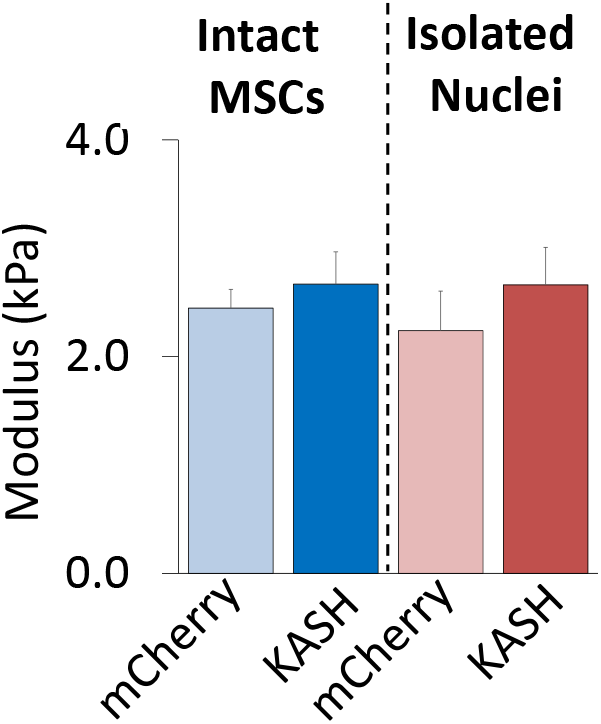
Compared to empty control plasmid (pCDH-EF1-MSC1-puro-mCherry), LINC-disrupted intact cells (N=10/grp) and isolated nuclei (N=20/grp) did not show any changes in stiffness. * p<.05, ** p<.01, *** p<.001, against control.

**Figure S4.**
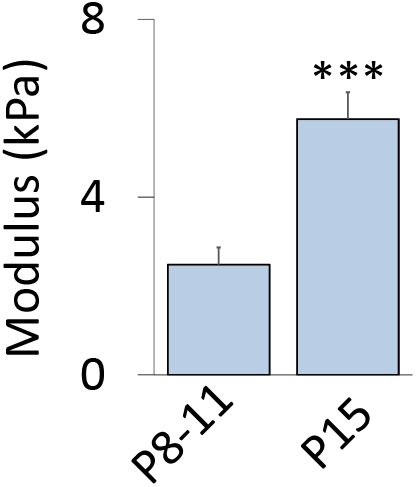
MSCs measured via AFM-based nanoindentation showed 132% higher elastic modulus in passage 15 samples (P15, N=10, p<.001) compared to samples between passages 8 and 11 (P8-11, N=45). * p<.05, ** p<.01, *** p<.001, against control or against each other.

**Figure S5.**
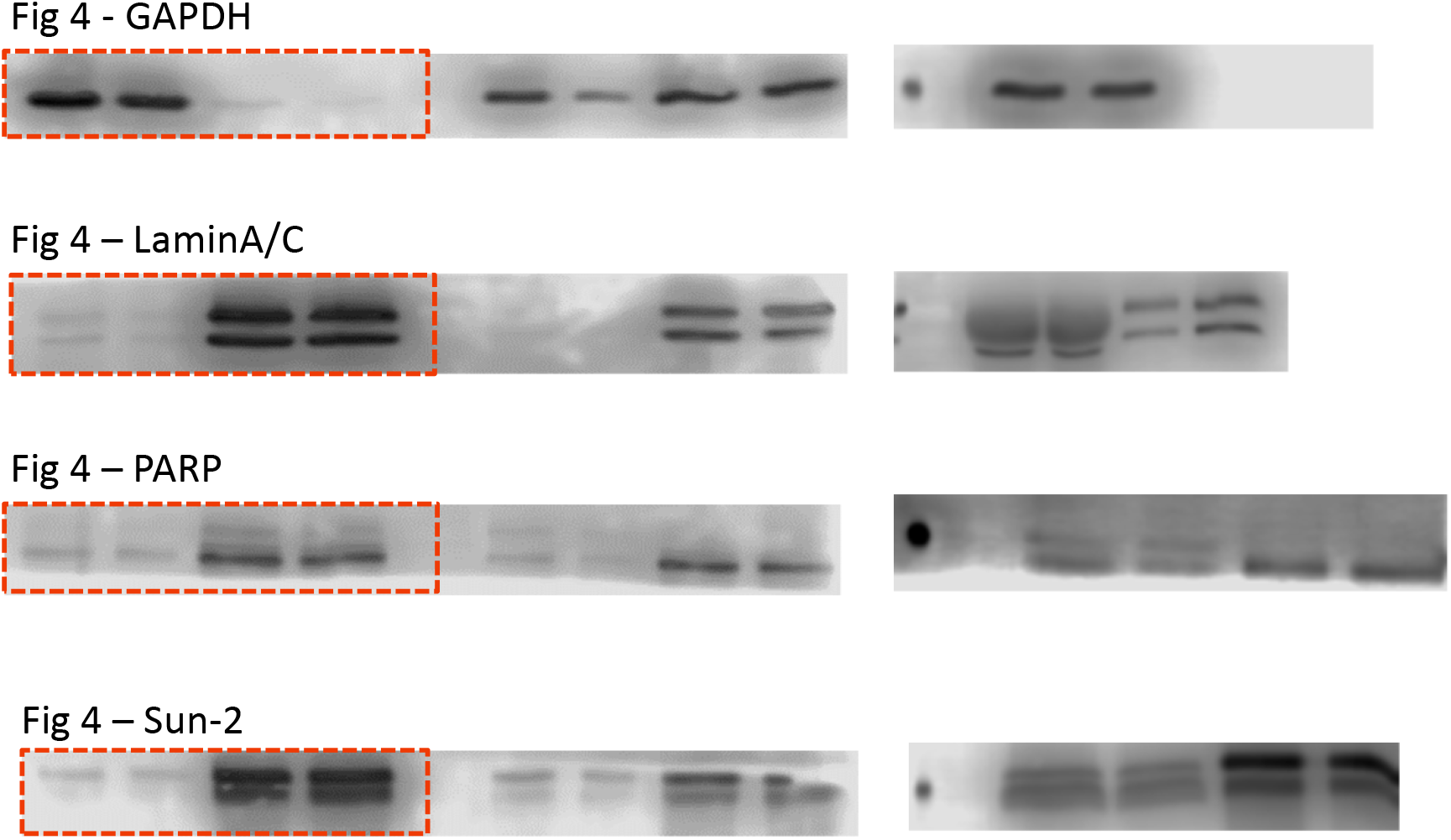
Unprocessed blots as obtained by LiCor C-DiGit blot scanner. Red lines outline the representative blots used.

### Supplementary Methods

#### Overexpression and Small Interfering RNA (siRNA)

PCDH-EF1-MCS1-puro-mCherry (mCherry control) and pCDH-EF1-MCS1-puro-mCherry-Nesprin-1αKASH plasmids were kindly provided by Dr. Lammerding. mdMSCs were transfected using 1μg DNA per 100,000 cells using LipoD293 transfection reagent (SignaGen Laboratories, Rockville, MD) according to the manufacturer’s instructions. 72 hr after the initial transfection, stably transfected cells were selected using 10 μg/ml puromycin. For transiently silencing specific genes, cells were transfected with gene-specific small interfering RNA (siRNA) or control siRNA (20 nM) using PepMute Plus transfection reagent (SignaGen Labs) according to the manufacturer’s instructions. The following siRNAs were used in this study: negative control for SUN-1 5′- GAAATCGAAGTACCTCGAGTGATAT -3′; SUN-1 5′- GAAAGGCTATGAATCCAGAGCTTAT- 3′; negative control for SUN-2 5′-CACCAGAGGCTAGAACTCTTACTCA-3′; SUN-2 5′- CAACAUCCCUCAUGGGCCUAUUGUG-3′. ′; negative control for LaminA/C 5′- UGGGAGUCGGAAGAAGACUCGAUCA-3′; LaminA/C 5′-UGGGAGAGGCUAAGAAGCAGCUUCA-3′.

#### Western Blotting

We used previously established western blot protocols (Uzer et al., 2018). Briefly, 20 μg of protein from each sample was separated on 9% polyacrylamide gels and transferred to polyvinylidene difluoride (PVDF) membranes. Membranes were blocked with 5% milk (w/v in TBST-T) and incubated overnight at 4°C with a specific primary antibody. Next, blots were washed and incubated with horseradish peroxidase-conjugated secondary antibody diluted at 1:5,000 (Cell Signaling) at room temperature for 1 hr. An ECL plus (Amersham Biosciences, Piscataway, NJ) kit and LiCor C-DiGit scanner were used to obtain quantitative data. Densitometry analysis was performed via NIH ImageJ using at least three independent experiments. During densitometry, each protein of interest was normalized to GAPDH or PARP, which were used as housekeeping proteins for whole cell or nuclear fractions, respectively. A list of primary antibodies used is given in **Table S2**.

#### Immunofluorescence and Image Analysis

Prior to experiments, nuclei were stained with Hoechst 33342 vital dye (Nucblue, ThermoFisher) according to the manufacturer instructions. Following the LIV protocol, intact MSCs or isolated live nuclei were fixed with 2% paraformaldehyde for 15 minutes. For chromatin intensity analysis, nuclei were imaged using an epifluorescence microscope (Revolve, Echo Labs). Chromatin intensity analyses were performed using a custom MATLAB script to select regions of nuclei. Hoechst stain was used to define the nuclear regions through the use of the blue channel. Each nucleus was individually defined, along with its intensity converted to 8-bit (i.e., a brightness range of 0-255). The mean brightness intensity of each nucleus was computed with heterochromatin being defined as +35 intensity relative to the average. The rest of the nuclear area was defined as non-heterochromatin, with the chromatin area and centroid then reported as output.

A heterochromatin area to nuclear area ratio was calculated to estimate average heterochromatin size as an indicator for changes in chromatin condensation. Individual heterochromatin regions were identified in isolated and intact nuclei for siRNA samples and vibrated samples using the previously described MATLAB technique. The ratio between heterochromatin average area and average nuclear area was computed and groups were compared using independent t-tests.

To quantify the nuclear geometry, the entire height of individual cells or nuclei were identified and further imaged using a Leica 6500 confocal microscope, which evenly divided each sample into sixteen vertical stacks **(Fig. S1**). Confocal image stacks were imported into FIJI ImageJ software (https://imagej.nih.gov/ij/). Using Hoechst 33342 as a landmark, nuclear height was quantified via counting the number of stacks between first and last slices with detectable, in-focus Hoechst 33342 signal using cross-sectional images. Next, the entire nuclear section was collapsed into a single image using “Average Intensity Projection.” Nuclear area was measured via tracing the outer circumference of Hoechst 33342. This allowed for the perimeter in each of the three planes of measurement (XY, XZ, and YZ) to be identified. From there, the “circularity” tool was used to compare each perimeter to that of a perfect circle. Averages for each plane of individual images were computed in Microsoft Excel for the two groups: intact MSC nuclei and isolated nuclei. The circularity values for each plane were then averaged to obtain a sphericity value for the two groups of nuclei.

### Supplementary Tables

**Table.S1:** Cell culture and pharmacological reagents

| Cell Culture and Pharmacological Reagents |  | Final Concentration |
| --- | --- | --- |
| IMDM | GIBCO | - |
| Fetal Calf Serum (FCS) | Atlanta Biologicals | 10% v/v |
| Penicillin/streptomycin | GIBCO | 1% v/v |
| SB415286 | Sigma Aldrich | 20μM |

**Table.S2:** Antibodies and concentrations

| Antibodies |  | Final Concentration |
| --- | --- | --- |
| PARP (9542T) | Cell Signaling | 1/1000 |
| GAPDH (5174S) | Cell Signaling | 1/1000 |
| LaminA/C (#4C11) | Cell Signaling | 1/1000 |
| Sun-2 (ab87036) | Abcam | 1/1000 |

**Table.S3:** Immunostaining antibodies, reagents, and concentrations

| Immunostaining Antibodies and Reagents |  | Final Concentration |
| --- | --- | --- |
| Alexa Flour 555 Donkey Anti-Rabbit | Invitrogen | 1/500 |
| Hoechst 33342 | Invitrogen | 1 μg/mL |
| LaminA/C (#4C11) | Cell Signaling | 1/300 |

**Table.S4:**
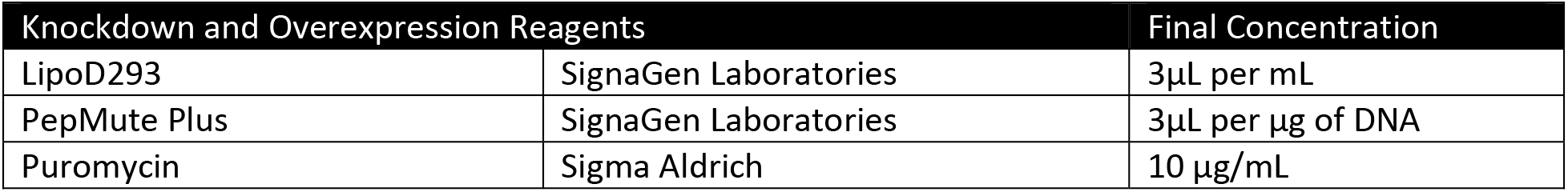
Overexpression reagents and concentrations

